# Recovery from fire affects spatial variability of nutrient availability in boreal aspen ecosystems

**DOI:** 10.1101/492389

**Authors:** S. Das Gupta, M.D. Mackenzie

**Affiliations:** Department of Renewable Resources, 348E South Academic Building, University of Alberta, Edmonton, AB Canada T6G 2G7; Canadian Forest Service, 2057 Northern Forestry Centre, Edmonton AB, Canada T6H 3S5

**Keywords:** Fire, Nutrient bioavailability, Spatial Variability, Aspen, Aboveground, Belowground, Boreal

## Abstract

Fire is a key driver of nutrient biogeochemistry in boreal ecosystems. Although a significant amount of research has been conducted to understand boreal fire ecology, it is still unclear how fire affects the spatial distribution of nutrients and what mechanisms are responsible for the post-fire recovery of spatial patterns. In this study, we examined spatial variability in soil nutrient bioavailability and related aboveground (AG) and belowground (BG) properties in three boreal aspen (*Populus tremuloides* Michx.) stands in northern Alberta at different stages of post-fire recovery. The studied sites include a 1-year old post fire stand (PF), a 9-year old stand at canopy closure (CC), and a 72-year old mature stand (MA). Ion exchange resin was used to measure nutrient bioavailability *in-situ* and was related to AG (vegetation and forest floor characteristics) and BG (soil microbial and chemical) properties. Significant spatial patterns were found in all three stands. PF stand had the greatest coarse scale spatial patterns (> 23 m) and availability of major macronutrients (N, P, and K). Shorter spatial range (5 to 10 m) of nutrient availability was observed in the stand with longest time since fire. Soil microbial activity was the strongest driver of nutrient availability in the PF stand, whereas contributions from aboveground variables such as understory vegetation, tree canopy cover, coarse woody debris (CWD), distance to nearest tree, and tree size was observed only in the CC and MA stands. The findings from the current study suggest that post-fire nutrient availability follows spatially predictable patterns, and confirm the hypothesis that stand replacing fire creates uniformity in nutrient availability and that the development of post-fire heterogeneity is a product of increasing ecosystem complexity.

## 1.0 Introduction

Wildfire is a major natural disturbance in upland boreal forests that causes abrupt changes in many ecosystems processes [1, 2]. The most distinct and immediate changes usually occur in vegetation structure and nutrient distribution in soils, both during and after fire events [3, 4]. Many pyrogenic ecosystems are fire adapted and crucial biogeochemical and community level processes depend on fire frequency and intensity [5, 6]. Besides influencing the key ecosystem processes such as nutrient cycling, gap dynamics and regeneration, fire alters the spatial variability of these processes [7], which eventually generates the commonly seen multi-scalar variability in ecosystem properties and creates a mosaic pattern on landscape. According to Odum, biogeochemical cycling of major nutrients becomes more spatially coupled and tighter as ecosystems progress towards maturity [8]. Several studies showed partial evidence in support of this hypothesis in different forested ecosystems [9, 10], however none explicitly tested the hypothesis from a spatial perspective, where space can be used as a surrogate for unmeasured variables [11] and spatial patterns can be used as an indicator of ecosystem complexity.

In recent years, a framework integrating different aboveground (AG) and belowground (BG) processes has been developed to better understand ecosystem functions and resilience [12, 13]. Although there is common agreement that the feedbacks between plant community and belowground organisms are strong drivers of both AG and BG processes such as competition and resource availability, our understanding of this relationship is still unclear due to the fact that these interactions are complex and happen at multiple spatial and temporal scales. In this study, we asked three specific questions on the development of spatial patterns in nutrient availability in boreal aspen ecosystem post-fire: (1) what spatial scale of variation do major macronutrients show in post-fire stands at different stages of recovery? (2) what degree of spatial association do these nutrients have with AG and BG properties? and (3) what AG and BG factors control the spatial patterns in nutrient availability in these stands?

High global variation or large-scale spatial structure in nutrient availability are expected in the PF stands (Question 1). Thus, we hypothesized that the semi-variograms will follow a linear or nugget model without any detectable spatial range and that spatial heterogeneity in nutrient availability would increase with time since fire (Figure 1). Several studies have indicated that the magnitude of nutrient heterogeneity decreases after stand replacing fire due to the homogenization of resources [14, 15] and the redistribution of ash [16]. However, development of fine-scale patchy structure in soil nutrients and microbial properties has also been observed one to ten years after fire [17, 18].

Spatial heterogeneity in stand structure and resources may increase with stand age due to competition, thinning and gap formation [19]. Aspen stands reach canopy closure quickly after fire, usually within 7 to 11 years, and significant changes take place in both AG and BG attributes related to nutrient availability at canopy closure [20, 21]. We speculated to see a fine to intermediate scale (~ 4 to 10 m) spatial pattern in nutrient availability in the CC stand due to tree canopy influences. A relatively large-scale (> tree scale; ~10 m) spatial pattern was expected for cation availability (Ca, Mg and K) at canopy closure due to their potential large-scale redistribution (e.g. via ash convection and transfer through forest floor) after severe fire, and relatively longer timeframe required to develop patchiness due to aboveground inputs, mostly through litter deposition [22, 23]. In MA stand, we expected to see a fine-scale (< 4 m) spatial pattern and distinct patchiness in nutrient availability due to an established microbes-root-canopy linkage.

For question 2, we hypothesized that spatial congruence between nutrient availability, AG and BG properties would increase with stand age due to an increased contribution from both microbial and tree level processes. The spatial association between AG and BG properties, and the degree at which all these three components (space, AG, and BG properties) are coupled together is termed here as spatial congruence. In boreal ecosystem, Lavoie and Mack (17) found increasing correlation between organic layer depth and soil properties with time since last fire, however their study did not account for spatial relationships in this correlation. Spatial coupling between AG and BG processes must increase over time as stands recover from fire disturbances. Plant traits have also significant influence on the development of spatial pattern in AG and BG properties. Differences in post-fire regeneration, growth, dispersal, and organic matter cycling between broadleaf and coniferous species control the differences in spatial patterns of BG properties in these stands to a large extent [24, 25]. For example, Sorenson, MacKenzie (26) in a reclaimed boreal forest showed nutrient availability in aspen stand is related to both canopy cover and forest floor mass, whereas only forest floor mass was associated to nutrient availability in pine stand. In a boreal coniferous forest Turner, Romme (27), however did not find spatial association between AG cover and N availability up to 4 years after wildfire and related this to a disjunct between the processes at the studied scale. Other evidences suggest that an increase in spatial association between vegetation and soil resources will occur with stand age as nutrient demands surpass supply as a result of tree competition [28, 29].

Finally, we expected to see a strong BG control on nutrient availability in the PF stand followed by stronger AG controls in the CC stand, and a joint AG-BG control in the MA stand (Question 3). Immediate post-fire patterns are likely to be associated with fire induced mortality of AG features which may take several years to re-establish [27, 30]. In boreal aspen ecosystems, understory vegetation and suckering species usually take over the AG space, but we believe their control on nutrient dynamics will not be evident immediately.

## 2.0 Materials and Methods

### 2.1 Study Sites

The study was conducted in the Athabasca oil sand region (AOSR), north of Fort McMurray, Alberta, Canada (56° 43′ N 111° 21′ W). The mean annual temperature in this region is 0.9°C and mean growing season (May – September) temperature is 13.3°C. Mean annual precipitation is 418.6 mm, of which 283.4 mm fall as rainfall during the growing seasons (Environment Canada 2014). Soils in the study areas are well to moderately-well drained Orthic Gray Luvisol (Typic Cryoboralf) with sandy loam to silty loam texture, and developed from till and glaciofluvial sediments [31]. Three boreal aspen stands representing variable stages of post-fire recovery were used in this study, viz. a one-year old post fire (PF) stand, a 9-year old stand at canopy closure phase (CC) and a mature 72-year old stand (MA). All three stands originated from mix-severity stand replacing fire. The most recent fire was in 2011 (Richardson Fire) which burned 576,000 ha and is the second largest documented fire in western Canada [32]. All the stands were on flat terrain with a very gentle slope (< 4%). Trembling aspen (*Populus tremulides* Michx.) was the most dominant tree species represented more than 95% of the basal area at all three sites and only a few sporadic white spruce (*Picea glauca* (Moench) Voss.) were found in the CC and MA stands. The PF stand had an aspen sucker density of 270, 000 stem ha^-1^, whereas the CC and MA stands had tree density of 1900 and 2150 stem ha^-1^, respectively. The maximum distance between the sites was 34 km. According to the ecosite classification of northern Alberta, all the three sites fall under the d1 ecosite phase (low-bush cranberry Aspen) [33]. More details on location, fire history and vegetation in the study area can be found in Das Gupta and Mackenzie (34).

### 2.2 Sampling Protocol

Sampling and field measurements were carried out in a 50 m × 20 m plot at all the three sites. A cyclic spatial sampling protocol with variable intervals [35] was used to capture both the scale and directionality in measured properties (Figure 2). In total 81 sampling points were established in 9 transects within a 1000 m^2^ plot which ensured a minimum detectable spatial lag of 0.5 m. Intervals between the sampling point along the transect were 0.5, 3, 6 and 9 m, and inter-transect intervals were 2 m and 4 m. The sampling orientation was reversed in the middle two transects to capture the anisotropy. All field measurements and sampling were done in May - August 2012.

### 2.3 Aboveground Properties

Aboveground properties were measured around each spatial point. Measured aboveground properties included tree location (XY coordinates), tree diameter (at breast height; DBH), number of aspen seedlings, canopy cover (%), understory vegetation cover (%; UV), bare ground (%; BGn), and coarse woody debris (% cover; CWD). Tree locations were only measured in the MA stand using a Nikon total station (Nikon DTM 352). Tree canopy cover (%) was measured using a convex densitometer, and percent cover of understory vegetation and coarse woody debris cover were measured using a 0.25 m^2^ square grid. Number of aspen seedlings was counted near each sampling point and 1 m^2^ sampling frame in the PF and CC stands. Basal area (BA) of nearest tree was calculated from tree DBH.

### 2.4 Belowground Properties

Belowground properties included forest floor depth (FD), root biomass, nutrient bioavailability, soil microbial and chemical properties. FD was measured at each spatial point by taking average of three measurements. Coarse root biomass (CRB) and fine root biomass (FRB) were calculated from the allometric equations developed by Chen, Zhang (36) and Brassard, Chen (37) (Appendix).

#### Nutrient bioavailability

Nutrient availability was measured using PRS^TM^ probes (plant root simulator probes; Western Ag Innovations Inc., Saskatoon, SK, Canada). These probes (cations and anions) have ion-exchange resin membranes which trap nutrients ions and nutrient supply rates are estimated based on the ion sinks adsorbed per surface area of membranes over the burial period [38]. Two pairs of cation and anion probes were vertically installed at the organic and mineral soil interface at each sampling point and left for 8 weeks to measure available nutrients under field conditions. Upon retrieval, probes were extracted with 0.5 M HCl and elutes were analyzed for nutrients. Ammonium (NH_4_^+^), nitrate (NO_3_^-^), and phosphate (PO_4_^2-^) were analyzed colourimetrically using a segmented flow Autoanalyzer III (Brand and Lubbe, Inc., Buffalo, NY.). Potassium (K^+^), sulfate (SO_4_^2-^), calcium (Ca^2+^) and magnesium (Mg^2+^) were quantified by inductively-coupled plasma (ICP) spectrophotometry (PerkinElmer Optima 3000-DV, PerkinElmer Inc., Shelton, CT). Nutrient availability was expressed as μg ion. 10 cm^-2^ 8 weeks^-1^.

#### Microbial Properties

For soil microbial and biochemical analyses, samples (organic layer + 5 cm mineral soil) were collected from each spatial point using a bulk density core. The core was surface sterilized and washed using ethanol (70%) in between samples to minimize any possible contamination and denaturation of enzyme products. Immediately after collection, soils were kept in a cooler with ice bags and brought back to the laboratory. Samples were then homogenized properly after carefully removing the coarse fragments (roots, twigs and stones). A sub-set of the samples were then stored at -20°C for extracellular enzyme activity and rest of the samples were stored at 4°C until further processing.

Soil microbial biomass C (MBC), N (MBN), and basal respiration (BR) were measured from an incubation experiment conducted using the soils stored at 4°C. Approximately 75 to 100 g soil was incubated for 10 days at 25°C in sealed Mason jar with alkali trap inside (0.5 M NaOH). Soil basal respiration was calculated from the trapped CO_2_ in alkali trap after titrating with HCl (0.5 M). Microbial biomass C and N was measured on the incubated samples using the fumigation extraction method [39]. Approximately 20 to 25 g soils were extracted in 0.5 M K_2_SO_4_ with a ratio of 1:2. Dissolved organic C (DOC) and N (DON) were measured on the unfumigated fractions of the soil extractions using Shimadzu TOC-V/TN analyzer (Shimadzu Corp., Kyoto, Japan).

Extra-cellular enzyme activity was measured on soils stored at – 20°C. Four enzymes were analyzed to quantify the potential extracellular enzyme activity responsible for C, N, P and S mineralization: (i) β-glucosidase (Bglu) (EC 3.2.1.21) responsible for breaking labile cellulose and other carbohydrate polymer chains, (ii) N-acetyle-β-D-glucosaminidase (NAGase) (EC 3.2.1.30), enzyme mainly catalyzes the hydrolysis of chitin, and convert it to amino sugars, which are major sources of N mineralization in soils [40], (iii) Phosphatase (Phos) (E.C. 3.1.3.2), phosphomonoesterases responsible for catalyzing the hydrolysis of esters and anhydrides of phosphoric acid [41], and (iv) Arylsulfatase (Sulf) (EC 3.1.6.1), enzyme that catalyzes hydrolysis of arylsulfate by breaking O-S bond and regulate mineralization of ester sulfate in soils [42]. Enzyme activities were measured using 4-methylumbelliferone (MUB; 10 mM) as a fluorimetric substrate. Details on the enzyme assay can be found in Sinsabaugh, Lauber (43) and Das Gupta, MacKenzie (44). Briefly, a 200 μl volume from a soil suspension (1 g soil homogenized in 100 ml 0.1M, pH 5 sodium acetate), and 50 μl of 200 μM substrate were pipetted into 96 well plates at 6 samples per plate. Microplates were incubated at 20° C in dark for 3 hours for β-glucosidase, N-acetyle-β-D-glucosaminidase and Arylsulfatase, and for 2 hours for phosphatase enzyme. Fluorescence was measured at 365 nm excitation and 460 nm emission, using a microplate spectrophotometer (Synergy HT, BioTek, Winooski, VT). Assay and control wells were replicated 8 times. Activity rates (μmol of converted substrate g^-1^ soil hour^-1^) were calculated on an oven dry mass (105°C) basis. Total C and N in soils were measured on ground samples using a Costech 4010 Elemental Analyzer System (Costech Analytical Technologies Inc., Valencia, CA, USA). Soil pH and EC were measured in 1:2 solution of deionized water using a pH electrode (Mettler Toledo EL20) and an EC meter, respectively.

### 2.6 Statistical Analyses

Coefficient of variation (CV) was used as an index of global variation in the measured variables. The CV is a simple metric of variability which can be interpreted easily and is comparable between different ecological properties [45]. Distribution patterns of nutrient availability, aboveground and belowground variables in different stands were compared in ordination space using principal component analysis (PCA). Multivariate comparisons among sites were done using multi-response permutation procedure (MRPP). PCA and MRPP analyses were done in PC-ORD ordination software [46].

Semi-variogram analysis was used to measure local variation i.e. spatial dependence of nutrient availability and other properties. Data were log transformed prior to analyzing semi-variogram and variogram modeling. Five variogram models viz. Linear, Gaussian, Exponential, Spherical and Nugget were tested to fit the empirical data. A combination of highest coefficient of determination (R^2^) and the lowest residual sums of square (RSS) was used to select the final model. Spatial dependence was calculated using the nugget coefficient, *n*_*c*_ which is a ratio of total variance (*c*_*0*_ + *c*_*1*_) and nugget variance (*c*_*0*_) [47].

Cross-variogram analysis was conducted between nutrients, soil and stand properties to check for any scale dependent spatial relationships. A positive cross-variance indicates spatial association whereas a negative variance means dissociation [48]. The spatial connections of AG and BG properties, and the degree at which all these three components (space, AG, and BG properties) are coupled together is termed here as spatial congruence. Spatial congruence among variables was measured by Kendall’s coefficient of concordance (W) [49, 50]. Variables were standardized and converted to a common scale between 0 and 100 before performing the test. Spatial congruence (W) is measured between 0 and 1, where, 0 means completely incongruent and 1 means perfect congruence. Spearman rank correlations between variables were also examined to know the strength and mode of bivariate relationships. Kendall’s W and Spearman rank correlations were done in in R v3.0.1 [51]. The ‘vegan’ package was used for calculating the Kendall’s W. Variograms were created using GS+ geostatistic software (V9.0, Gammadesign software). Spatial autoregressive (SAR) models were developed between nutrients and environmental variables, and contribution of space was estimated using two SAR model viz. spatial error model and spatial lag model [52, 53]. SAR analysis was done in R [51] and Geoda, an R based open source geospatial software [54]. Details on the semi-variogram modeling, SAR calculation, and interpretations are given in the Appendix.

## 3.0 Results

The PF stand was characterized by an open canopy, greater aspen sucker density, bare ground cover, and dissolved organic N compared to the CC and MA stands. Canopy covers in the CC and MA stands were 74% and 93%, respectively. A number of soil microbial properties including MBC, Bglu and NAGase activities, and C:N ratio were similar in the PF and CC stands. The MA stand had overall greater soil microbial biomass and activity than the PF and CC stands (Appendix Table 1).

**Table 1.** Mean, coefficient of variation (CV), and variogram model parameters for available macronutrients (μg 10 cm^-2^ 8 weeks^-1^) in three boreal aspen stands at different stages of post-fire recovery in northern Alberta.

| Stands | Nutrients | Mean (SE) | CV (%) | Range (m) | Spatial dependence | Model | R<sup>2</sup> |
| --- | --- | --- | --- | --- | --- | --- | --- |
| Post fire | TIN | 7.25 ± 1.07 | 133 | - | - | NUG | - |
| | P | 31.7 ± 2.89 | 82.0 | > 23 | 0.24 | LIN | 0.14 |
| | S | 36.07 ± 2.49 | 62.2 | 6.5 | 0.81 | GAU | 0.40 |
| | K | 344.4 ± 17.8 | 46.7 | - | - | NUG | - |
| | Ca | 1216 ± 61.4 | 45.4 | 1.1 | 0.88 | GAU | 0.12 |
| | Mg | 90.43 ± 4.38 | 43.6 | - | - | NUG | - |
| Canopy closure | TIN | 3.34 ± 0.24 | 63.7 | 8.0 | 0.50 | GAU | 0.17 |
| | P | 5.74 ± 0.40 | 62.8 | 3.1 | 0.83 | SPH | 0.18 |
| | S | 25.9 ± 2.88 | 98.7 | > 23 | 0.36 | LIN | 0.25 |
| | K | 177.5 ± 13.6 | 68.8 | > 23 | 0.52 | GAU | 0.48 |
| | Ca | 1326.1 ± 64.6 | 43.8 | > 23 | 0.38 | LIN | 0.28 |
| | Mg | 144.3 ± 5.2 | 32.2 | - | - | NUG | - |
| Mature stand | TIN | 3.96 ± 0.16 | 36.4 | 9.8 | 0.60 | SPH | 0.39 |
| | P | 6.64 ± 0.50 | 68.1 | 5.8 | 0.51 | SPH | 0.15 |
| | S | 54.3 ± 5.1 | 83.7 | 10 | 0.61 | SPH | 0.40 |
| | K | 250.5 ± 17.1 | 61.6 | > 23 | 0.50 | LIN | 0.21 |
| | Ca | 1079.4 ± 34.7 | 28.9 | > 23 | 0.53 | EXP | 0.26 |
| | Mg | 308.5 ± 9.24 | 27.9 | 17.6 | 0.51 | GAU | 0.46 |

### 3.1 Nutrient Availability

Significant differences were found in nutrient profile among the three fire affected stands. Total inorganic nitrogen (NH_4_^+^ and NO_3_^-^), P and K availability were the highest and Mg availability was the lowest in the PF stand. The CC stand had the lowest S availability followed by the PF and MA stand. Nitrate was the main form of N in all the three stands and the availability was found highest in the PF stand (4.09 ± 2.45 μg 10 cm^-2^ 8 weeks^-1^) followed by the CC (2.18 ± 1.26 μg 10 cm^-2^ 8 weeks^-1^) and MF (2.37 ± 1.30 μg 10 cm^-2^ 8 weeks^-1^) stand. Likewise, P availability in the PF stand was 31.7 (± 26) followed by 5.67 (± 3.67) in the CC and 6.84 (± 4.84) μg 10 cm^-2^ in the MF stand (Table 1).

Distribution of nutrient availability, soil microbial properties and stand characteristics in different stands are shown in the PCA ordination plots (Figure 3). The ordination axes explained 20 to 57% of variation in the AG and BG properties. MRPP analysis indicates significant differences among the three stands for all the three ecosystem matrices. The most distinct differences were observed for AG properties, whereas nutrient availability and BG properties had common distribution pattern in the three stands, yet significantly different than each other.

### 3.2 Spatial Variability

Higher global variation (CV) in N, P, Ca and Mg availability was found in the PF stand than the other stands which gradually decreased over time with the lowest in the MF stand. For example, CV of available TIN was 133.17, 63.64 and 37.37% in the PF, CC and MF stand, respectively. On the other hand, lower global variation was also observed in the PF stand than the older stands for certain other nutrients such as S (62.18%) and K (46.68%) (Table 1).

Most of the nutrients in the PF stand either had large-scale or no spatial pattern, except for S and Ca. Nutrients in the CC and MA stand, however, showed spatially structured patterns. Sulfur and Ca availability had strong spatial dependency in the PF stand (≤ 6.5 m), which showed large-scale dependency (≥ 21 m) in the CC and MF stand. Nitrogen and P availability were both spatially structured at < 10 m in the CC and MF stand but had either a pure nugget model or large-scale structure in the PF stand (Table 1 and Figure 4).

### 3.3 Spatial Association

Spatial associations between nutrients, AG, and BG properties were measured using cross-variance analysis (Figure 5). Nitrogen, P, and S in the PF stand had spatial association with enzyme activities at ≤ 10 m scale, but no aboveground association was detected for these nutrients (Figure 5; a, f and i). The CC and MA stands, on the other hand, showed spatial association of these nutrients with both enzymes, substrates (DOC), and aboveground variables such as understory vegetation cover and tree distance, although nutrient-enzyme association was only detected in the CC stand (Figure 5; b and j). Nitrogen availability showed a negative spatial association with UV at 7 m scale in the CC stand and a positive spatial association with tree distance at 9 m scale in the MA stand. The spatial control from understory vegetation and forest floor depth on the availability Mg and K was also found only in these stands, except Mg had a fine-scale (2.4 m) negative association with understory vegetation in the PF stand (Figure 5; l, m and n).

Number of significant correlations between nutrients, aboveground and belowground is presented in Appendix (Table 2). Out of 126 possible correlations (6 nutrients x 21 variables), the PF stand had only 18 significant cases (14%), the CC stand had 20 significant cases (16%), and the MA stand had 28 significant cases (22%). Similar trend was also found when only belowground resources such as DOC, DON, MBC, MBN, C, N, and FD (proxy of organic matter) were considered. Out of such 42 possible correlations (6 nutrients x 7 resources), PF stand had only 6 significant cases (14%), which was 8 (19%) and 11 (27%) in the CC and MA stands, respectively. Number of significant correlations between nutrients was, however, the highest in the PF stand (66%; 10 out of 15 correlations) than the CC (27%; 4 out of 15) and MA (40%; 6 out of 15) stands (Table 2). Correlations between nutrients in the PF stand were all positive, whereas, the relationships changed in most cases in the CC and MA stands. For example, TIN showed a negative relationship with S and a positive relationship with P in the PF stand, but the reverse was true in the CC and MA stands.

Spatial congruence (as measured by Kendall’s W) between nutrient availability, AG and BG properties (AG-BG-NUT) was the lowest in the PF stand which gradually increased with stand age (Figure 6). Spatial congruence was 0.28, 0.33, and 0.40 in the PF, CC and MA stand, respectively. Congruence between only AG and BG properties (AG-BG) was also greater in the CC and MA stands than the PF stand. Available nutrients, however, had the strongest congruence in the PF stand followed by the CC and MA stands.

### 3.4 Aboveground-Belowground Controls

Spatial autoregressive models (Error and Lag) were used to account for the aboveground and belowground contributions to nutrient availability in different stands. A clear influence of microbial processes, especially enzyme activity, was found in the models of the PF stand. Nitrogen was the only nutrient influenced by an aboveground variable (percent bare ground). Phosphorus and Ca appeared to be influencing the availability of most of the macronutrients in the PF stand. On the other hand, both aboveground and belowground properties exerted significant influence on nutrient availability in the CC and MA stands. Significant belowground variables in the spatial models were DOC, MBCN, BR and enzyme activities. Different sets of aboveground variables appeared to have significant control on nutrients in these two stands. For example, CWD and UV were significant consistent variables in most of the spatial models in CC stand, whereas, Tdist., BA and UV were consistent in the nutrient models of MA stand. Significant spatial structure in the macronutrient regression models (SAR terms) was detected only for Ca in the PF, for K and Ca in the CC stand, and for all other nutrients except Ca and K in the MA stand (Table 3).

## 4.0 Discussion

### 4.1 Spatial Variability of Soil Nutrients

This study presents evidence of high nutrient availability and low spatial variability immediately after fire in boreal aspen ecosystems. Although post-fire ecosystem properties are generally assumed to be heterogeneous [14, 55, 56], there are only a few studies that have actually looked at spatial relationships and their dynamics from an aboveground and belowground perspective, and a lot of these studies were done in coniferous stands, with a few in boreal ecosystems. We hypothesized that post-fire nutrient availability would be high, with a high variation (CV) and less spatial predictability. This was true in the PF stand and spatial heterogeneity increased with stand development through CC and into MA. The findings are also in line with our expectation to observe a nugget or linear semi-variogram models in the PF stand and more spherical models in the MA stand due to sharp edge formation in the nutrient patterns (Figure 1).

Fire effects on soil nutrients have been studied extensively in different ecosystems. A significant number of studies have reported an increase in mineral nutrients in post-fire environment [57–59]. In a comprehensive review, Certini (60) reported that fire usually increases the long-term availability of nutrients especially N and P, whereas the increase in Ca, Mg and K availability is relatively ephemeral. Effect of recent burn on major elements and plant available nutrients in boreal aspen stand is scarce [61]. In boreal forest ecosystems, Neff, Harden (62) did not find any significant fire effect on P, Ca, Mg and K stocks which they attributed to the post-fire erosional loss and chemical immobilization due to the increased availability of Al and Fe. In an experimental burn study in boreal Alaska, Harden, Neff (3) found higher N, P, Ca, Mg and K in the burned soil which corroborates with our findings except for Mg.

The rate at which these nutrients recover to a pre-fire level depends on the ecosystem type, fire severity and vegetation at the time of fire [61]. Similar nutrient availability in the CC and MA stands in the current study might suggest that ecosystem recovery from wildfire disturbance can happen at a faster pace, i.e. within 10 years of disturbance. Belowground properties of these two sites also showed major distribution overlap as indicated in the ordination plot (Fig. 3c), which further support this assumption. A variable timeframe of post-fire recovery in nutrient conditions is reported in the literature with a maximum of 35 years [5, 63]. Soil microbial properties, on the other hand, has been shown to recover within 12 years after fire [64]. Adams and Boyle (65) reported an ephemeral increase in P, Ca, K and Mg in a mixed aspen stand one month after fire which decreased to pre-fire level within 5 months. The initial soil heterogeneity in post-fire environment is assumed to be created by spatial variability in fire severity and the understory vegetation that survived from fire [55, 66]. Ephemeral post-fire heterogeneity in major plant available nutrients may become homogenized quickly due to uniform abiotic conditions (e.g. soil temperature and moisture, and light availability) and vigorous natural regeneration of clonal species such as aspen in the current study [67].

Spatial information on the availability of macronutrients other than N is limited in pyrogenic boreal ecosystems. Variability in the methods used for measuring nutrient availability in different studies is another constraint that makes such comparisons difficult. Smithwick, Mack (18) reported a spatial pattern of N mineralization and organic C at 8.3 m and 2.5 m scales, respectively in a 1-year post-fire boreal black spruce dominated stand. The multiple scale dependency of N and C availability in their study was attributed to topographical and microclimatic variation. Lavoie and Mack (17), however, did not find any spatial structure in N mineralization and C pools (TC, C:N and BR) in a 8-year old boreal black spruce-aspen stand, but spatial heterogeneity gradually appeared in the older stands of their studied post-fire chronosequence, which corroborates with the current study. In a dry tropical forest, Hirobe, Tokuchi (68) also found a similar fire effect on N mineralization as in our study. The spatial range of N mineralization decreased from ≥ 9 m to 3.2 m with time since last fire. They also found strong spatial dependence in soil properties in the stand with longest time since fire (35 years). Dietrich and MacKenzie (69) also reported a no to very weak spatial dependency of essential macronutrients (N, P, K, and S) in a post-fire (1-year old) boreal jack pine stand. In an oak-aspen forest Adams and Boyle (65) found that irrespective of fire intensity and variation due to coarse woody debris on site, the fire induced nutrient availability was fairly uniform and did not show any significant differences among treatments. However, they found significant differences in cation concentrations (Mg and K) in sub-soil layers shortly after fire and attributed this variation to the accelerated mineralization of combusted organic matter and reduced plant uptake.

We did not find any detectable spatial pattern in K and Mg availability, but a very fine-scale pattern in Ca availability in the PF stand. Large-scale patterns were detected for these nutrients in the CC and MA stands. The missing spatial pattern in base cations in the post-fire environment might be related to leaching loss and absence of canopy input to the forest floor layer. Tree canopy and forest floor play significant role in base cation cycling through microclimatic modification, litter input, and throughfall [70]. The fine-scale spatial pattern in Ca availability in the PF stand could be related to the increase in soil acidity from the high nitrification rate as usually observed immediately after fire and harvesting events [71]. The observed large-scale spatial patterns in base cations in the CC and MA stands can be attributed to the development of canopy structure and forest floor which generally occurs at stand scale. Bengtson, Basiliko (72) also reported very large-scale or no patterns in cation availability in a mature coniferous forest in coastal British Columbia, which they think were generated due to landform factors such as topography and might not be related to the spatial variability in organic and mineral soil horizons. Spatial variability in cations can also originate from geological variability (i.e. variability in chemical composition of minerals) and variable weathering rate of minerals [73].

### 4.2 Spatial Relationships with Soil Nutrients

Disturbances which dramatically alter the biogeochemical and other abiotic fluxes (e.g. fire) can significantly change the spatial connections between the AG and BG factors [49]. We hypothesized that the post-fire stand would have a weak spatial congruence between AG and BG properties which we expected to increase with recovery from fire. Spatial congruence as measured by Kendall’s W and correlation between AG and BG properties in different stands supported our hypothesis. However, post-fire nutrient availability showed significant correlation which was not observed in the other two stands (Table 2). The strong positive correlations between macronutrients in the PF stand indicate a clear effect of fire on the availability of these nutrients, which could be through ash convection or downward movement of nutrients via stemflow, charring, and sloughing off plant parts [59]. The missing relationship between N and other nutrients further indicates the high demand for this nutrient in post-fire environment (Table 2). Different interactions among nutrients in the CC and MA stands point out that the observed relationship in the PF stand is ephemeral and might not have originated from spatial regulatory processes. Hart, DeLuca (74) mentioned a strong link between post-fire plant functional groups that are spatially more determinative for structuring belowground biotic community and so the related processes such as nutrient availability. Large intermittent disturbance like stand replacing wildfire can change plant functional traits and the spatial associations between belowground processes by killing most of the dominant tree species and creating areas with homogenous microclimatic conditions [24]. In fire adapted ecosystems such as the boreal forests, the broken spatial link between aboveground and belowground properties can recover quickly through the vigorous regeneration and development of forest floor, as the current study shows. Several other studies reported a positive correlation among nutrients after fire. For example, Chorover, Vitousek (22) found a strong correlation between anions and cations in forest floor and soil solutions collected four months after fire in a mixed conifer forest in California. Ion concentrations in mineral soil solution showed greater sensitivity to fire than forest floor. The high cation availability post fire decreased with time which they attributed to plant uptake and downward movement into soil profile. Murphy, Johnson (75), however, reported non-significant differences in base cations between burned and unburned plots in a mixed coniferous forest stand one year after fire and identified inherent site variability as the principal mechanism.

Since the aboveground vegetation structure in the PF stand was yet to develop, a strong correlation in nutrient availability can be interpreted as indirect fire-induced effect and not of spatial origin. The lack of spatial autocorrelations in some of these nutrients in the PF stand and gradual increase of spatial association (Kendall’s W and cross-variograms) in AG-BG and AG-BG-Nutrients in the CC and MA stands (Figure 5 and 6) also supports this. Study by Lavoie and Mack (17) corroborates with the higher number of significant correlations among resources in the mature stands in our study. In their study, the number of significant correlations between AG and BG properties increased from 3 in the youngest post-fire site to 17 in the oldest site of their boreal fire chronosequence. Hirobe, Tokuchi (68) reported a gradual increase in the strength of correlation between N mineralization and soil moisture content in a dry tropical forest chronosequence. However, such relationships between AG and BG properties in disturbed ecosystems may not follow a consistent pattern throughout the stand recovery process due to environmental and seasonal resource gradients and may show spatial associations at multiple scales [18, 76].

### 4.3 Aboveground-Belowground Controls on Nutrient Bioavailability

Although there are detail studies on nutrient availability in post-fire boreal ecosystems, to our knowledge this is the first study which accounted for both aboveground and belowground controls on nutrient bioavailability using spatial models. The findings indicate a belowground biochemical control on nutrient availability in the PF stand with almost no spatial structure, and a joint aboveground-belowground control in the CC and MA stands with significant spatial structures. Substrate availability (DOC) and C mineralization efficiency (qCO_2_) are driving the availability of most of the macronutrients in the CC stands, whereas substrate quality (C:N) appeared as the most consistent belowground driver of nutrients in the MA stand. On the other hand, extracellular enzyme activities were the important controlling factors for the availability of almost all the macronutrients in the PF stand (Table 3). The gradual increase in AG contribution to overall nutrient availability and spatial congruence with stand age suggest a joint AG-BG control in the stands past canopy closure phase. Significant space term in the regression models of nutrient availability in the MA (mostly) and other stands indicates a possibility of nested spatial patterns at smaller scales which were not captured by the current sampling protocol or important plant and soil variables were overlooked.

Strong enzymatic control on nutrient availability in the PF stand might be due to the presence of partially decomposed and thermally altered OM in this stand. Fire often creates a flush of mixed molecular weight compounds in soil which may in turn trigger variable microbial activity in different soil layers [77]. Taş, Prestat (78) reported higher carbohydrate-degrading enzyme activity in top soils (0 – 10 cm) of post-fire boreal forest in Alaska and together with OM characterization using ^13^C-NMR they showed that a preferential decomposition of labile C and preservation of hydrophobic alkyl C pools exist in post-fire soils despite a large reduction of C content and fresh C input. The energetics of OM decomposition dictates microbes to first work on the labile, low molecular weight substrate and then shift towards high molecular weight and more persistent compounds [79, 80]. We did not find any significant enzymatic control on N availability in the PF stand. But a strong DON control indicates that there is still a flush of labile N and microbes are probably not utilizing N from other sources. Fire can preferentially favor chitonolytic organisms (mainly bacteria) to produce more chitin in the post-fire environment [81]. In high N environment, as in the PF stand, chitin can depress chitinase (NAG) activity [82]. A positive correlation between N and NAG in the PF stand, however, indicates that such enzymatic relationship might be of importance for N acquisition in post-fire systems. A negative relationship between P and phosphatase, and S and sulfatase indicate an end-product inhibition. A recent study by the authors [44] found similar relationships in a reclaimed aspen stands and suggested P limitation and S abundance (from atmospheric deposition) as driving mechanisms.

The CC and MA stands showed significant aboveground control on nutrient availability which was expected as a result of biomass accumulation, forest floor and canopy development, and changes in forest composition [83]. In a fire chronosequence of coniferous forests in Montana, MacKenzie, DeLuca (84) showed that forest floor thickness and aboveground biomass gradually increased with time since fire following a log-linear pattern, and significantly influenced the speciation of available N along the trajectory of secondary succession. Their results also indicated a higher N mineralization rate during the early stages of succession and a lower mineralization rate during the later stages, which were attributed to the difference in organic matter quality (i.e. more labile during early succession vs more persistent at later stages). In boreal forest ecosystems, DeLuca, Nilsson (85) also reported similar findings. Interestingly, the aboveground drivers of nutrient availability were not the same in the CC and MA stands. Stand level attributes such as forest floor thickness, understory vegetation and coarse woody debris cover were mostly significant in the CC stand, whereas more individual tree-based attributes such as distance to nearest tree, tree basal area and canopy cover were mostly significant in the MA stand.

## 5.0 Conclusions

Nutrient availability in pyrogenic environment often evolves through complex multi-scalar interactions, which might determine the mosaic pattern of vegetation and biogeochemistry in mature landscape; however, such relationships and their spatial variability depend very much on the mode of disturbance and ecosystem types. Pyrogenic boreal aspen forests are some of the least studied ecosystems in terms of the development in spatial heterogeneity in biogeochemical properties and associated mechanisms. The findings from this study support some of the previously proposed disturbance hypotheses by [86] and present some evidence of AG-BG controls on nutrient bioavailability at different stages of post-fire recovery in aspen mixedwood boreal forests. In general, the findings suggest that spatial variability in AG and BG properties must be considered when considering nutrient availability in post-fire ecosystems. Although the post-fire site had a very high number of aspen suckers, spatial variability in nutrient availability in this site does not seem to be controlled by their high density; rather soil microbial and enzymatic properties were important. This suggest that the spatial patterns in tree seedling of clonal species such as aspen may not be driven by the variability in nutrients in post-fire stands, at least during the early post-fire recovery phase. A synergistic control of both AG and BG variables on nutrient availability at the canopy closure and mature stands may explain the spatial structuring of vegetation and individual tree driven processes (e.g. organic matter cycling) in these stands.

The spatial information (range and variability) and predictive models of nutrients as developed in the current study are useful for studies that are designed to compare changes in ecosystem properties before and after large-scale natural disturbance such as wildfire. Of special interest to ecosystem reconstruction after resource mining, this study will work as natural benchmark and will guide the reclamation efforts to create a target ecosystem with specific spatial patterns in nutrient availability, aboveground and belowground properties. We, however, acknowledge that the observed patterns in the current study are based on single stand spatial dynamics and may not be extrapolated beyond the scale of the study.

## Acknowledgements

Financial support for this study came from the Environmental Research and Reclamation Group (ERRG) of the Canadian Oilsands Network for Research and Development, and the National Science and Engineering Research Council of Canada. We thank Pak Chow, Megan Lewis, Mark Howell, Sawyer Desaulniers, and Nicole Filipow for their help in the field and laboratory.

